# Assessing visual performance during intense luminance changes in virtual reality

**DOI:** 10.1101/2024.04.16.589684

**Authors:** Niklas Domdei, Yannick Sauer, Brian Hecox, Alexander Neugebauer, Siegfried Wahl

## Abstract

During indoor-outdoor transitions humans encounter luminance changes beyond the functional range of the photoreceptors, leaving the individual at risk of overlooking harmful low-contrast objects until adaptation processes re-enable optimal vision. To study human visual performance during intense luminance changes, we propose a virtual reality based testbed. After linearization of the headset’s luminance output, detection times were recorded for ten participants. The small (FWHM = 0.6 degree) low-contrast stimuli appeared randomly in one of four corners (±10 degree) after luminance changes of three magnitudes within 1 or 3 seconds. Significantly decreased detection times were observed for the conditions with simulated self-tinting lenses compared to lenses with fixed transmission rates after luminance decreases. In cases of luminance increases all detection times were similar. In conclusion, the proposed virtual reality testbed allows for studying vision during or after steep luminance changes and helps to design technical aids like self-tinting lenses.

**Highlights:** The HTC Vive headset provides a practicable luminance range of 3 magnitudes, allowing the simulation of indoor-outdoor transitions.

Visual performance after steep luminance decreases was significantly better with simulated self-tinting spectacles reducing the encountered luminance step.

Different transmission rates of the simulated lenses did not affect the detection times for small low-contrast stimuli after luminance increases.

## Introduction

The human visual system is functional across an enormous luminance range of about 14 magnitudes.^1^ For example, humans can observe a sunny outdoor scene, but also detect individual photons.^2,3^ This is possible, first, because of two different photoreceptor cell types in the human retina: the cones and the rods. The rods are operational at scotopic and mesopic light levels, while the cones are functional at high light levels (mesopic and photopic) and enable color vision.^4^ Second, adaptation plays an important role. Without adaptation, the individual photoreceptor has a functional range of about 3 magnitudes. The overall photoreceptor activation to luminance relationship follows a sigmoid function and has a central part of 1.5 log10 units, where luminance is linearly correlated with activation level.^5– 7^ This observation matches closely with the luminance range within a natural scene, spanning approximately 2 magnitudes.^1,8^

However, transitions between indoor and outdoor, for example when driving into a tunnel or walking into a building on a sunny day, put the photoreceptor’s functional range at its limits, with luminance changes of up to 3 magnitudes.^9^ Thus, to enable optimal vision and to perceive low-contrast objects or obstacles, a shift of the photoreceptor’s functional range is indispensable and is achieved via adaptation.^10^ A rapid (hundreds of milliseconds) adaptation mechanism of the visual system is the adjustment of the pupil diameter. Theoretically, a range of 2 to 8 mm is possible, thereby allowing adjustment of retinal illuminance by a factor of 16 (or 1.2 log10 units).^11^ Practically, this range only occurs in healthy young adults during transitions from photopic to scotopic illuminance. In the scenario of a young driver entering a tunnel, pupil diameter changes from 4 to 7 mm were reported,^12^ which changes retinal illuminance by a factor of about 3 (or 0.5 log10 units). For elderly people (>60 years) pupil diameters are decreased, but the range is similar.^12^ At the same time, the photoreceptors adapt as well, but this process is rather slow compared to the pupil response: For example, to shift the cone’s functional range by 1 magnitude, it takes about 30 seconds.^10^

Taken together, this leaves a significant timescale, within which humans are unable to perceive low-contrast objects, resulting in increased visual discomfort during indoor-outdoor transitions like tunnel entrances in daylight.^13^ In the case of tunnels, this situation is tackled in modern tunnels by sophisticated lighting.^14^ Additionally, to ease the visual discomfort elicited from indoor-outdoor transitions, for example when entering a building on a sunny day, the solution can be photochromic self-tinting lenses.^15,16^ However, photochromic lenses have limited usability, as they depend on ultraviolet light and reaction times on average of approximately 210 seconds to switch between states at room temperature,^17^ which is too slow for most of the encountered indoor-outdoor transitions. The latest solutions are ultraviolet-light independent electrochromic lenses,^18,19^ which have highly variable characteristics regarding their switching times and transmission values. To this point it is unclear which electrochromic lens characteristics (e.g., the available transmission range, minimum and maximum transmission, or response time), are physiologically relevant or in case of a trade-off more important to support the visual system by preventing the need for adaptation.

We propose a virtual-reality (VR) testbed to simulate the encountered changes in illumination during indoor-outdoor transitions to evaluate any electrochromic lens characteristics regarding its capability to reduce visual discomfort. For an objective read-out of such an assessment, a visual performance test is implemented as well, emphasizing the degree of helpfulness of an electrochromic lens with the given characteristics.

## Results

### Linearization

Linearization of the used HTC Vive VR headset’s output luminance was achieved by using a shader to directly apply an achromatic 8bit RGB value (in the following referred to as grey value) to a surface covering the entire display. The use of a custom shader ensured full control of the surface’s ultimately rendered grey value. Firstly, at each grey value (in the range 0 to 255), the resulting light power was measured repeatedly three times while continuously increasing or decreasing grey values (see Figure 1). Repeated power measurements showed a relative standard deviation of about 2 % for grey values between 17 and 255. For low grey values (grey values between 8 and 16), the relative measurement variability was about 5 % and for the residual lowest grey values (0 to 7) measurement variability was about 20 %. Secondly, at 255, 0, and 4 intermediate grey levels the luminance was measured, confirming a direct relationship between power meter readings and actual luminance (see Table 1). Repeated luminance measurements showed a variability of about 3 %. At grey value 0 a residual power of approximately 13 nW and luminance of 0.014 cd/m^2^ were measured on average. At 255 the power was about 93 µW and luminance was 140 cd/m^2^ yielding a maximum contrast ratio of approximately 1:10,000 or a possible luminance range of 4 magnitudes.

**Table 1:** Comparative listing of the HMD’s average output power and luminance measurements.

| Grey value | Power meter reading [$\mu W$] | Log10 (power) | Contrast ratio | Luminance [$cd/m^2$] | Log10 (lum.) | Contrast ratio |
| --- | --- | --- | --- | --- | --- | --- |
| 255 | 93 | 1.97 | 1.0 | 140 | 2.15 | 1.0 |
| 192 | 48 | 1.68 | 1.9 | 76 | 1.88 | 1.8 |
| 128 | 19 | 1.29 | 4.8 | 30 | 1.48 | 4.7 |
| 64 | 4.3 | 0.63 | 22 | 6.6 | 0.82 | 21 |
| 10 | 0.095 | -1.02 | 979 | 0.14 | -0.85 | 1000 |
| 0 | 0.013 | -1.89 | 7152 | 0.013 | -1.89 | 10769 |

**Figure 1:**
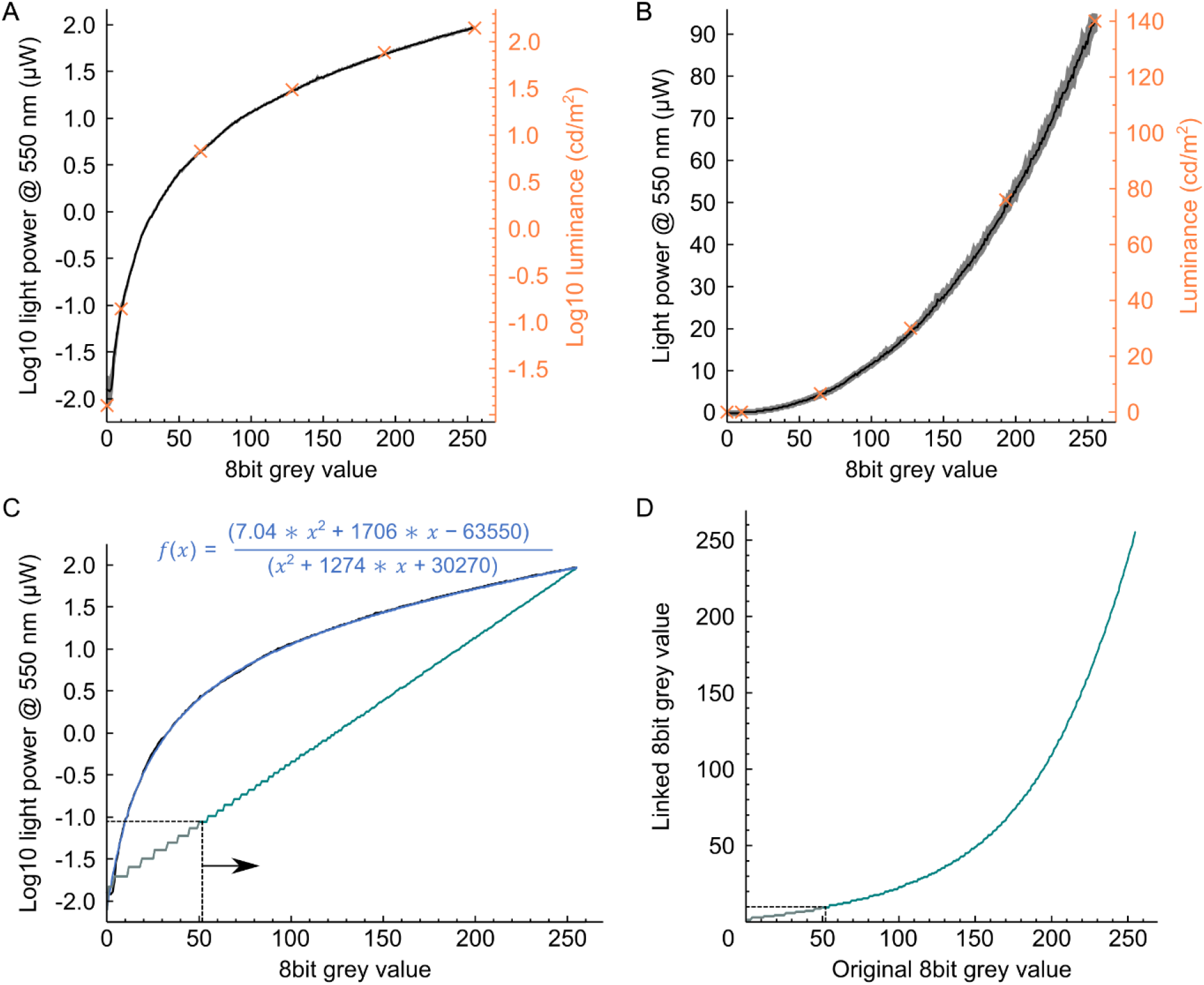
Output power linearization for the HTC Vive HMD. A & B) Set grey level and corresponding average logarithmic or linear scaled output power in µW. The grey shaded area shows ± 1 standard deviation of repeated measurements. Luminance was measured at selected data points (orange crosses), confirming the validity of the used power values for linearization. C) The average logarithmic power values were fitted with a rational function (blue, equation stated). 8bit grey values were linked according to D). From the linearized output (turquoise line in C), only 8bit grey values ≥10 were used leaving a residual luminance range of 3 magnitudes.

The relationship between grey values and their average output power was fitted with a rational function (Figure 1C):

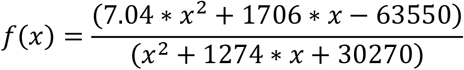

Using this fit, linearization was achieved by creating a look-up-table linking each grey value between 0 and 255 to a new grey value (Figure 1D) enabling equal changes in luminance on a logarithmic scale. The logarithmic scale was chosen due to “Fechner-Law” stating that the perception of luminance scales proportionally to logarithmic changes, meaning that for a sensation of linear luminance changes, logarithmic changes are required.^20^ In Unity, the linearization was realized based on the “Dictionary” class to link the respective grey value pairs in a look-up-table. For the final transition simulation, grey values from 10 to 255 were used, because of the high measurement uncertainty for low grey values and relatively large luminance increments between single steps for grey values <10. According to Table 1, the lower grey value of 10 corresponds to a luminance of 0.14 cd/m^2^, while the upper grey value of 255 corresponds to 140 cd/m^2^. Thus, the selected grey value range enabled a range of 3 magnitudes for luminance changes.

### Transmission implementation

This look-up-table was also used for the calculations to convert any required transmission, of a currently simulated spectacle lens, into the respective alpha value. To this end, it was mandatory to set the project’s color space to “gamma”.^21^ At the beginning of each frame update the luminance value behind the lens “Lum_behind_” is calculated as the product of the current wall luminance “Lum_original_” and the transmission of the simulated lens:

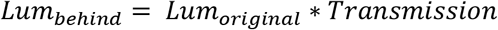

The respective 8bit grey value (“8bitGrey_behind_”) was determined by using the look-up-table in reverse (method “FirstOrDefault”) and finding the closest entry for the needed modified luminance “Lum_behind_”. The simulated lens’ alpha could then be calculated as

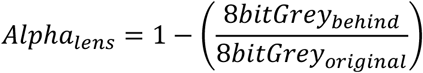

and was applied to the canvas simulating the lens.

While normal spectacle lenses or sunglasses have a static transmission, electrochromic or photochromic lenses are designed to have a dynamic transmission. This dynamic transmission was implemented with a linear as well as a more realistic exponential switching behavior. The simpler linear switching allowed a direct and therefore easier control of maximum and minimum transmission, and response time. The more complex, but realistic exponential switching was implemented by using MATLAB to compute an exponential function according to the desired specification. Here, two different electrochromic lenses were simulated. Both had a similar range of transmission states, with a maximum transmission of 90 % and a minimum transmission of 10 %. The “fast” switching electrochromic lens had a response time (defined by reaching 90 % of the target transmission) of about 1 sec, and the “slow” switching had a response time of approximately 3 seconds (see Figure 2A). The resulting exponential functions describing the switching from clear to dark were:

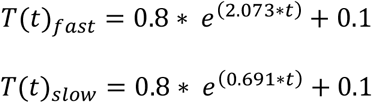

And for switching from dark to clear:

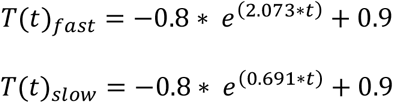

In the following psychophysical assessment procedure for testing visual performance, transmission changes of electrochromic lenses were automatically triggered by a virtual binary light sensor with a threshold of 4 cd/m^2^ (8bit grey = 54). Between the light sensor switch and the onset of the transmission, a simulated electronic delay of 100 msec was added (Figure 2B).

**Figure 2:**
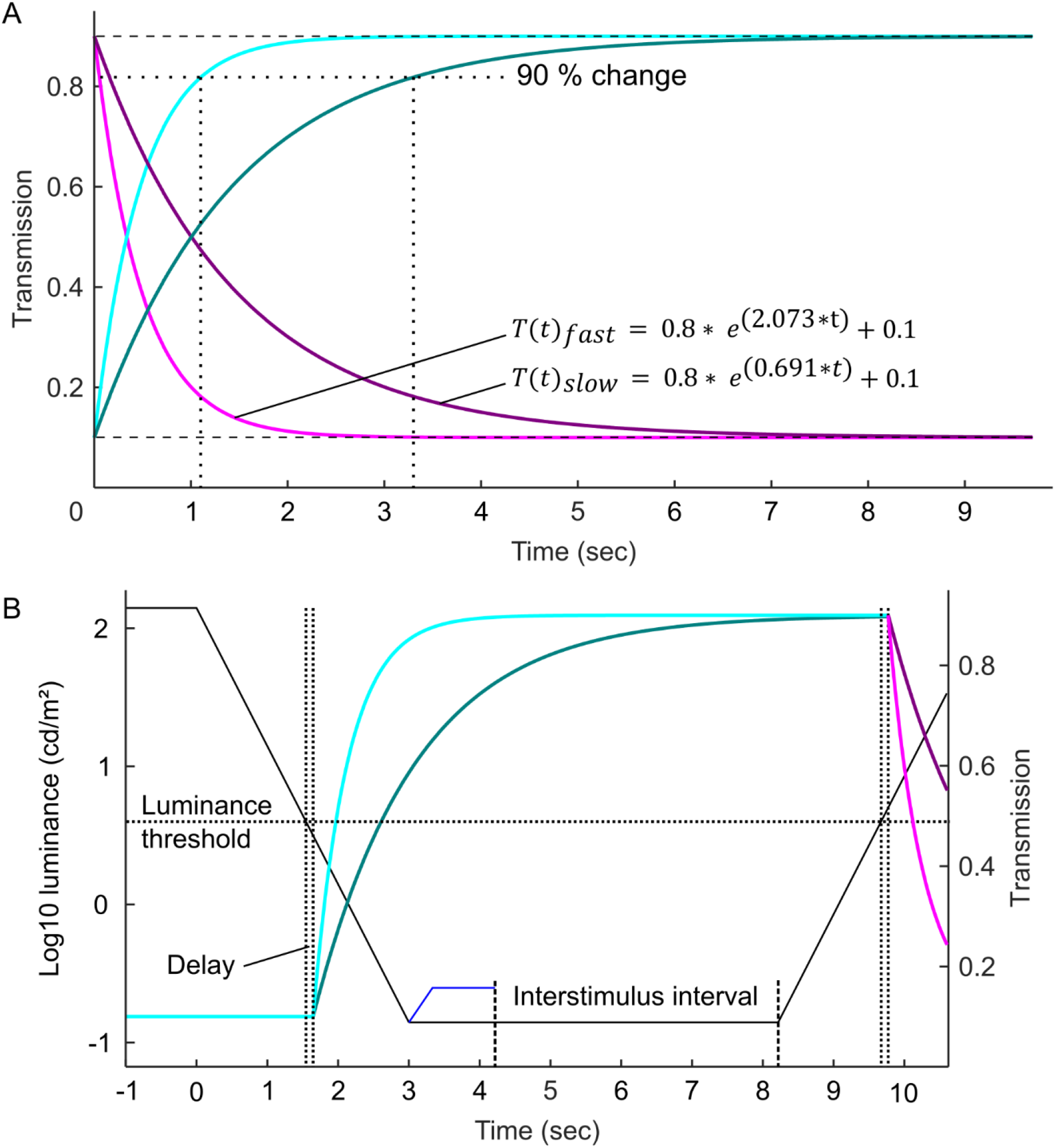
The electrochromic lenses’ transmission change over time is given by an exponential function. A) Transmission changes for the fast and slow electrochromic. The respective response times, given by a 90% completed change (dotted lines), are about 1 and 3 sec. B) Complete illustration of the electrochromic assessment protocol. Each trial starts with a luminance change across several magnitudes. Crossing the luminance threshold triggers the transmission change after a set delay (here 100 msec). The stimulus fades in when the target luminance is reached and is deleted on detection. Before the next trial starts, the interstimulus interval allows the visual system to adapt to the new lighting.

### Psychophysical assessment

The psychophysical assessment procedure of visual performance consisted of 40 trials for each of the 8 different conditions (see Table 2), with increasing and decreasing lighting scenarios tested consecutively. Because of the logarithmic linearization, the luminance could be changed at a constant rate of 1 log10 units per second (Figure 2B) or 3 log10 units/sec. When the target luminance was reached the stimulus faded in at 0.3 log10 units /sec. The starting luminance of the stimulus was adjusted to ensure a fixed fade-in time of 333 msec. The target luminance was empirically determined to make sure that stimuli were only just perceivable, resulting in unequal Weber-contrast values for the sunglasses mode (2.5) compared to normal or electrochromic lenses (0.7) and between luminance increases and decreases (see Table 3 and 4). Weber-contrast was similar across conditions with about -0.2 at high luminance. The stimulus appeared randomly in one of four corners (Figure 3).

**Table 2:** Test scenarios for psychophysical assessment of visual performance with and without electrochromic (eChromic) lenses. Each scenario included 40 trials in total (20 trials for room lights switching off and 20 trials for switching on). A detailed description of each lens type is given in the main text.

|  |  |  |  |
| --- | --- | --- | --- |
| eChromic_fast (1 sec)<br>Fast light (3 log10 units /sec) | eChromic_slow (3 sec)<br>Fast light | Normal<br>Fast light | Sunglasses<br>Fast light |
| eChromic_fast (1 sec)<br>Slow light (1 log10 units /sec) | eChromic_slow (3 sec)<br>Slow light | Normal<br>Slow light | Sunglasses<br>Slow light |

**Table 3:**
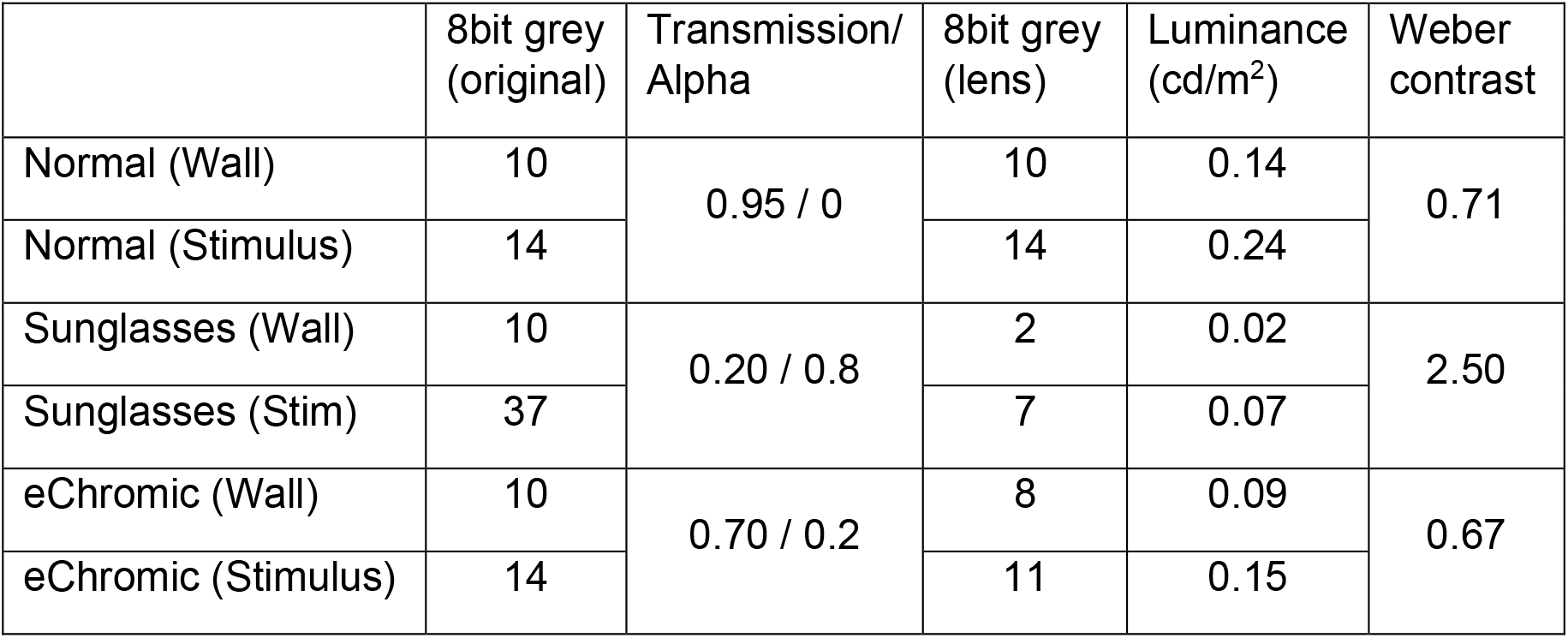
Weber contrast for stimuli at low light conditions.

| | 8bit grey (original) | Transmission/ Alpha | 8bit grey (lens) | Luminance ($cd/m^2$) | Weber contrast |
| --- | --- | --- | --- | --- | --- |
| Normal (Wall) | 10 | 0.95 / 0 | 10 | 0.14 | 0.71 |
| Normal (Stimulus) | 14 |  | 14 | 0.24 |  |
| Sunglasses (Wall) | 10 | 0.20 / 0.8 | 2 | 0.02 | 2.50 |
| Sunglasses (Stim) | 37 |  | 7 | 0.07 |  |
| eChromic (Wall) | 10 | 0.70 / 0.2 | 8 | 0.09 | 0.67 |
| eChromic (Stimulus) | 14 |  | 11 | 0.15 |  |

**Table 4:**
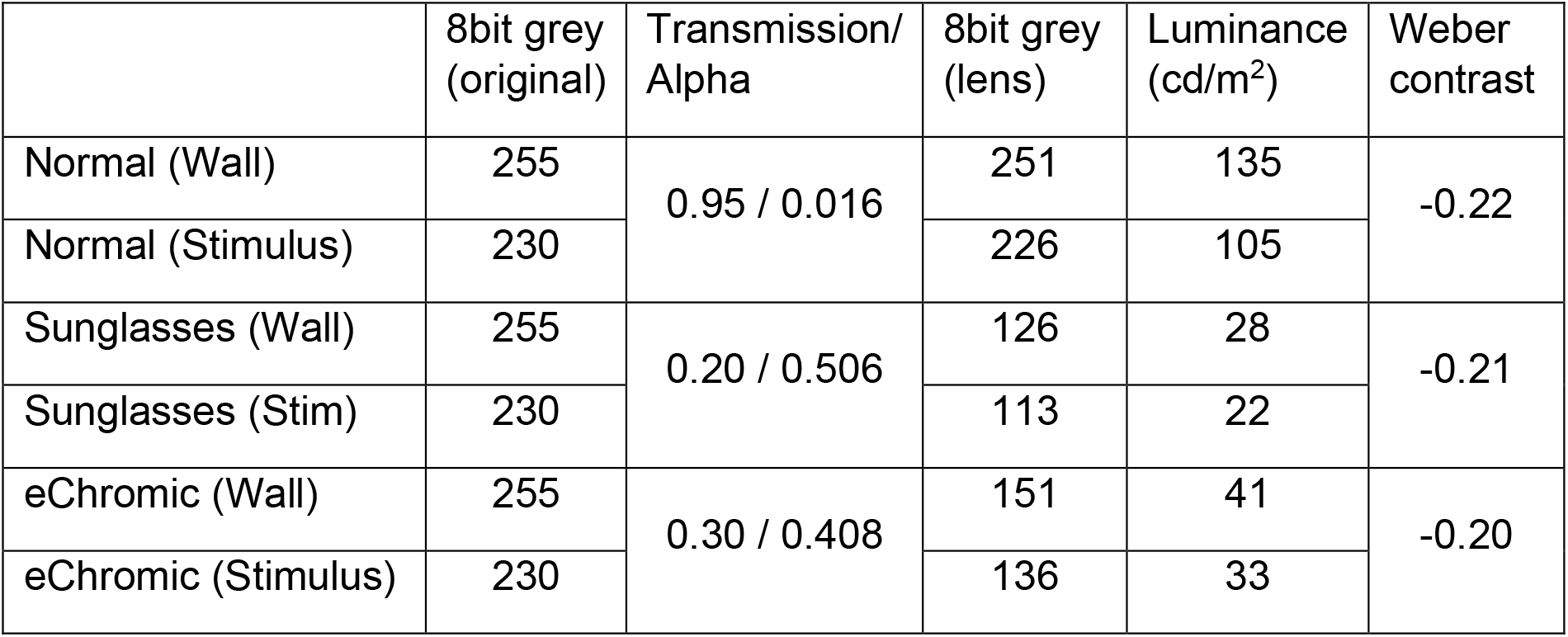
Weber contrast for stimuli at bright light conditions.

|  | 8bit grey<br>(original) | Transmission/<br>Alpha | 8bit grey<br>(lens) | Luminance<br>(cd/m<sup>2</sup>) | Weber<br>contrast |
| --- | --- | --- | --- | --- | --- |
| Normal (Wall) | 255 | 0.95 / 0.016 | 251 | 135 | -0.22 |
| Normal (Stimulus) | 230 |  | 226 | 105 |  |
| Sunglasses (Wall) | 255 | 0.20 / 0.506 | 126 | 28 | -0.21 |
| Sunglasses (Stim) | 230 |  | 113 | 22 |  |
| eChromic (Wall) | 255 | 0.30 / 0.408 | 151 | 41 | -0.20 |
| eChromic (Stimulus) | 230 |  | 136 | 33 |  |

**Figure 3:**
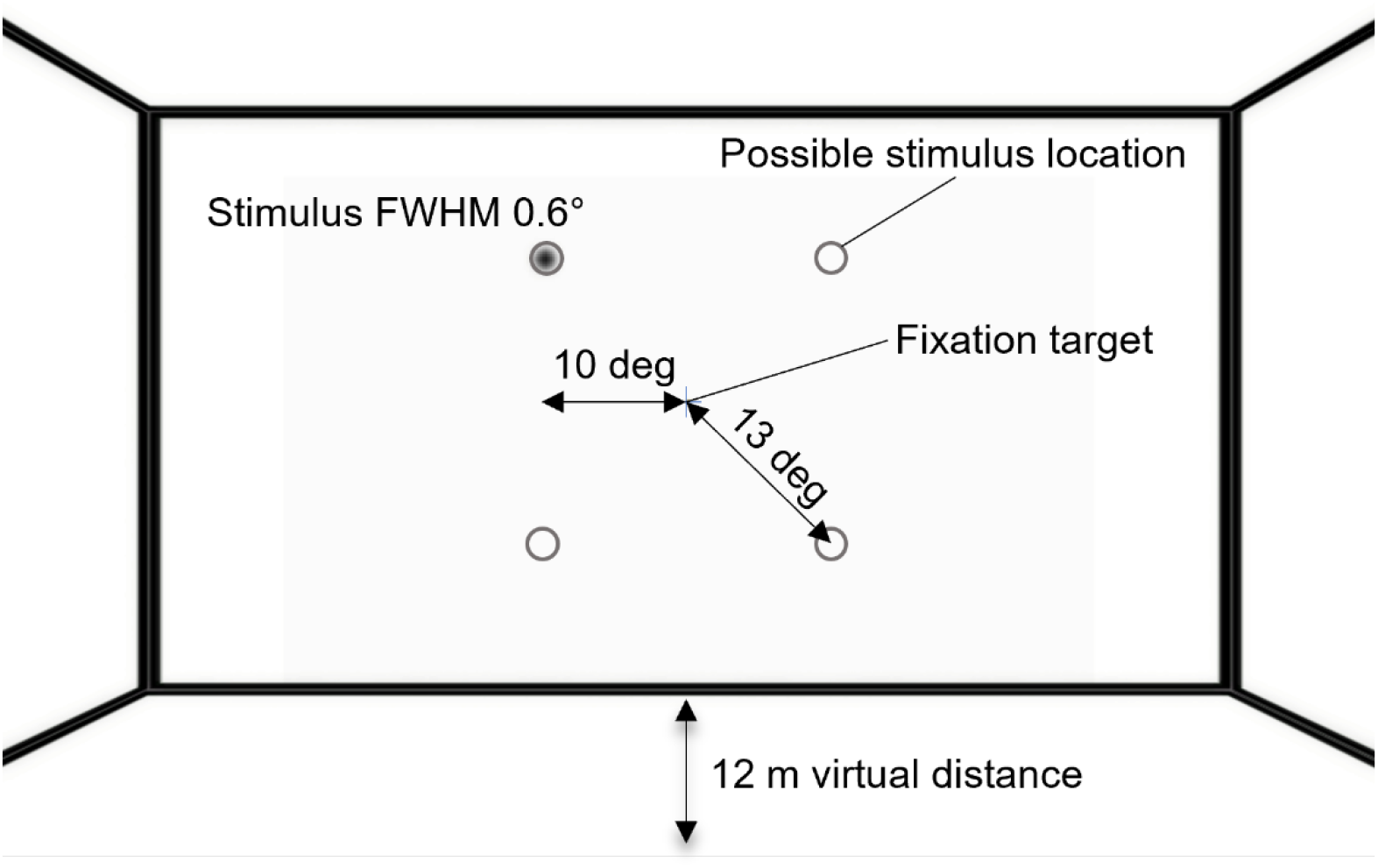
VR room design of the testbed with the corresponding stimulus parameters for the visual performance assessment.

Participants (see STAR methods for details) pressed the corresponding key on the numpad on stimulus detection, for example, “7” for the upper left corner. When the location was correctly reported, the stimulus object was deleted. Before the next trial was initialized an interstimulus interval of 4 seconds had to be awaited allowing the visual system to adapt to the new light condition. In the following, detection times are stated as Median and interquartile range (IQR) in seconds. The statistical details are given in the STAR methods.

After a quick luminance decrease, detection times were significantly lower in the fast electrochromic condition (1.20 (0.30), all p<0.01; Figure 4A). Detection times increased for the slow electrochromic lens (1.62 (0.19)) but were significantly lower (all p<0.01) than in normal (2.04 (0.36)) or sunglasses mode (2.59 (0.79)). The observed increase in detection times for sunglasses compared to normal lenses was significant as well (p<0.05). After a slow luminance decrease, again detection times were lowest for the fast electrochromic lens (0.91 (0.25); Figure 4B), followed by the slow electrochromic lens (1.02 (0.18)). This difference in detection times between the two electrochromic lens modes was not significant. However, significantly higher detection times were observed for the non-electrochromic lens modes (all p<0.01). For normal and sunglasses lens mode detection times were similar and not significantly different (1.32 (0.30) and 1.31 (0.80), respectively). When luminance was increased by 3 magnitudes detection times were similar across different lens conditions and the two different change rates (for 3 log10 units/sec: 1.15 (0.16), 1.10 (0.20), 1.09 (0.17), 1.19 (0.15); for 1 log units / sec: 1.20 (0.22), 1.09 (0.15), 1.06 (0.17), 1.12 (0.23) for fast and slow electrochromic, normal and sunglasses respectively; Figure 4C and D). Detection times were significantly higher for simulated sunglasses compared to normal lenses after quick luminance increases (p<0.05) and significantly higher for the fast electrochromic lenses compared to all other lens modes after slow light increases (p<0.01 and p<0.05).

**Figure 4:**
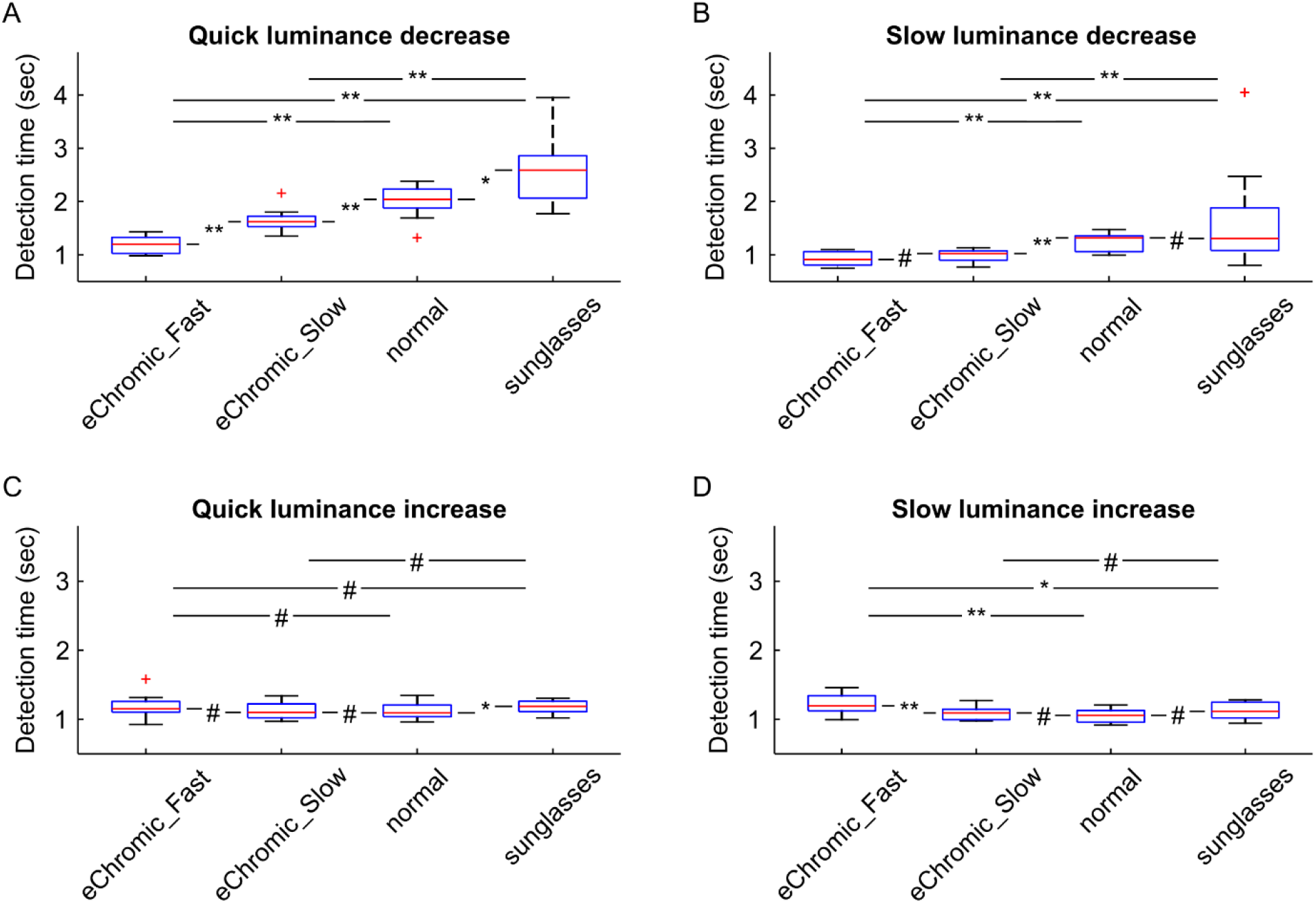
Detection times after luminance changes for all different lens conditions. A) and B) show detection times after quick (3 log10 units/sec) and slow (1 log10 unit/sec) luminance decreases. C) and D) show detection times after luminance increases. Asterisks denote significance level (# p>0.05; * p<0.05; ** p<0.01; N=10)

## Discussion

Using an HTC Vive virtual reality headset, we were able to set up a testbed to study human visual perception during and after large luminance changes, which are encountered in our daily life during indoor-outdoor transitions like driving into a tunnel where luminance changes by 3 magnitudes. Because the functional range of photoreceptors can only cover changes of 1.5 log10 units, adaptation processes are required to shift the photoreceptor’s sensitivity accordingly. During this adaptation period, humans are unable to perceive low-contrast objects, like a small obstacle, which could be a tripping hazard. Therefore, indoor-outdoor transitions elicit a high level of visual discomfort, which can also be objectively assessed by measuring fluctuations in pupil diameter.^13^ Modern technical aids e.g., electrochromic spectacles, may provide an option to support the visual system during large luminance steps by changing their transmission and thereby reducing the amount of luminance change. However, the required characteristics of an electrochromic spectacle to make for a sufficient aid are unclear. To this end, a simulation of transmission changing spectacle lenses was added to the indoor-outdoor transition testbed, as well.

While the headset could not display the absolute luminance values of outdoor scenes, the available luminance range after linearization of the headset’s luminance output enabled us to mimic luminance changes of indoor-outdoor transitions across 3 magnitudes. This range is sufficient to simulate the luminance changes encountered when driving into a tunnel, where the luminance changes from 5000 cd/m^2^ outside to 4 cd/m^2^ inside the tunnel.^9,22^ When entering a building on a sunny day the required range for a proper simulation is a little bit smaller as the median illuminance drops by 2 magnitudes from about 4 to 2 log10 units inside.^23,24^

Detection times observed here, were similar across the different conditions after luminance increases (on average 1.1 sec). However, after luminance decreases, detection times were significantly different between the tested conditions. Trials with simulated electrochromic lenses had the lowest detection times of about 1.2 seconds on average across conditions followed by normal static transmission lenses (2.0 and 1.3 seconds for quick and slow luminance changes, respectively) and simulated sunglasses (2.6 and 1.3). These observed detection times are much higher compared to simple reaction times of about 400 msec reported for peripheral testing with mesopic luminance.^25,26^ There are several reasons that could explain the large difference between our study and previous reports. First, the stimulus used was a blob with a full width at half maximum (FWHM) of 0.6 degrees and thereby up to three times smaller than the 1.5 to 2 degree diameter circular stimuli used in other psychophysical experiments.^22,25,26^ Second, being a blob, the stimulus did not have a distinct grey level step at the edges, to make detection more challenging. Third, the stimulus did not suddenly appear but contrast increased during the first 330 msec of the presentation time to make detection even more challenging. These specific stimulus characteristics were chosen empirically, to emphasize differences in visual performance for the tested conditions.

Because stimulus presentation time was not limited to a short presentation time and participants were allowed to look freely, the stimulus location on the retina was unclear. However, simple reaction times between central and peripheral stimuli (7.5 degree eccentricity, 0.8 degree diameter) are similar at about 300 msec.^27^

Due to the removal of any temporal or spatial sharp onsets, the task implemented here is more related to a search task than a simple reaction time experiment. This is also reflected in the fact that when neglecting the 330 msec fade-in time, the observed detection times are close to reported search reaction times of about 750 msec for a small set size.^28^

Interestingly, detection times were similar in the electrochromic lens condition for in- or decreasing luminance, however, the stimulus contrast was different (about -0.2 and 0.7 for luminance increases and decreases, respectively). This observation is in accordance with previous studies, reporting increasing simple reaction times for decreasing background luminance but the same Weber-contrast of small peripheral (10 degree eccentricity) stimuli.^25,26,29^ For example, to achieve similar simple reaction times for 1 cd/m^2^ and 0.1 cd/m^2^ background luminance, the Weber contrast needed to be significantly increased from 0.2 to 0.7.^25^ At a background luminance of 0.01 cd/m^2^, which is comparable to the sunglasses condition in our experiment (see Table 3), even at the highest contrast setting of 0.8, reaction times were 100 msec slower than for a background luminance of 0.1 cd/m^2^. It can only be assumed from the presented data, that a stimulus contrast of > 3 would be required to achieve reaction times comparable with the 0.1 cd/m^2^ background luminance and 0.7 contrast condition.^25^ Which again bears resemblance to the study here, where stimulus contrast for the sunglasses condition was much higher (2.5) than for the other two conditions, for similar detection times between the static transmission scenarios. Additionally, for a simple visual discrimination task under low mesopic viewing conditions it was reported that visual performance in terms of d-prime, a measure of sensitivity, decreases, too.^30^ As a side note, these observations provide a scientific justification why class 4 sunglasses (transmission rate <8%) are not suitable and forbidden for driving: Inside a tunnel light level would be diminished to a mesopic luminance resulting in significantly increased reaction times for spotting a potential hazard.

While detection times were slightly elevated for luminance decreases compared to luminance increases depending on the tested condition, at the same time, detection times were similar across all conditions for luminance increases. An explanation for this might be given by the observation that adaptation to light is significantly faster than dark adaptation.^31^ Because dark adaptation relies on the metabolism of the visual cycle replenishing the photopigment, the gap between light and dark adaptation grows with increasing age while light adaptation stays on the same level across ages.^31^ Therefore, elderly people would benefit even further from electrochromic spectacles reducing the need for adaptation to see well during and shortly after an indoor-outdoor transition.

For future studies utilizing this testbed, it could be worthwhile to fade in the stimulus by increasing its size rather than increasing contrast. In this way, stimulus emergence would be closer to a real-world scenario where e.g., a driver moves closer to an obstacle. Furthermore, including pupil tracking could help to provide an objective readout for the participant’s visual discomfort during rapid luminance changes, as it was shown that visual discomfort correlates with fluctuations in pupil diameter.^13^ Such pupil diameter fluctuations were reported for an increased cognitive load as well.^32,33^

With the available range of 4 log10 units, the HTC Vive headset could be utilized for dark adaptometry, if equipped with neutral density filters. Dark adaptometry can be useful as an early indicator of e.g., age-related macular degeneration^34^ or other retinal diseases related to systemic vitamin A deficiency, where impaired dark adaptation leads to a complete dark-adapted rod sensitivity.^35,36^

### Limitations

In the shown proof-of-concept psychophysical experiment detection times for small low-contrast stimuli after steep luminance changes with a rate of 1 or 3 log10 units /sec were applied. These rates are much steeper compared to entering a tunnel with modern lighting where luminance changes stepwise by 3 log10 units within 7 seconds (about 0.4 log10 units /sec).^9,22^ However, to our best knowledge no literature exists on the timescale of luminance change rates when walking into a building or driving into a tunnel without modern lighting.

## Conclusions

The presented testbed to study human visual perception related to large changes in luminance demonstrated that the available luminance output range of the used HTC Vive headset sufficiently covers the typically encountered range of real-world luminance changes. The observed detection times across the different conditions tested here are in good accordance with previous studies reporting increasing reaction times for low background luminance levels at the same stimulus contrast or similar reaction times when stimulus contrast was increased for low background luminance levels. The implementation of a self-tinting lens simulation into this testbed will be useful to determine the required range and time scale of transmission changes to sufficiently bolster visual perception during indoor-outdoor transitions. Future studies assessing electrochromic lens characteristics may include the usefulness of intermediate transmission states as well as the user acceptance of manual or automatic transmission control.

## Author contributions

Conceptualization: N.D. and S.W.; Methodology: N.D.; Software: N.D., Y.S., B.H., and A.N.; Investigation: N.D.; Writing – Original Draft: N.D.; Writing – Review & Editing: All authors; Visualization: N.D.; Funding Acquisition: S.W.; Resources: S.W.; Supervision: S.W.

## Acknowledgements

Funding: The study was supported by the German Research Foundation (DFG) under SFB 1233, Robust Vision: Inference Principles and Neural Mechanisms, TP TRA, with project number 276693517. Open Access funding enabled and organized by Projekt DEAL.

The authors thank all participants for their contribution in this study.

## Data and code availability

Data from this experimental study have been deposited at Mendeley and are publicly available as of the date of publication. The DOI is listed in the key resources table.

All original code has been deposited at github and is publicly available as of the date of publication. The URL is listed in the key resources table.

Any additional information required to reanalyze the data reported in this paper is available from the lead contact upon request.

## STAR Methods

### Lead contact

Further information and requests for resources should be directed to and will be fulfilled by the lead contact, Siegfried Wahl.

### Experimental model and study participant details

#### Study participants

For the psychophysical evaluation procedure, a total of 10 healthy participants (4 males; mean age 29 ± 4 years) were recruited in this study, all members of the workgroup at the University of Tübingen. All participants had normal or corrected-to-normal vision, with no known retinal pathologies.

The study adhered to the tenets of the Declaration of Helsinki. The ethics authorization to perform the measurements was granted by the Medicine Faculty Human Research Ethics Committee from the University of Tübingen. Prior to data collection, the experiment was explained in detail to the participants, and written informed consent was collected from each participant. All data were pseudonymized and stored in full compliance with the principles of the Data Protection Act GDPR 2016/679 of the European Union.

#### Method details

#### Hardware

Based on a preliminary study from our lab, it was decided to use the HTC Vive (HTC Cooperation, Taoyuan, Taiwan) VR head-mounted display (HMD) as it provides the largest range of luminance levels known for HMDs,^37^ due to its organic light-emitting diode (OLED) display technology. The individually specific output light power levels of the used HMD were assessed with a power meter (818-SL/DB sensor with 843-R power meter, Newport, USA) and a self-written Unity3D (version 2021.3.12f1; Unity Technologies, CA, USA) environment setting the HMD’s OLED displays to a given achromatic grey level between 0 and 255. For the linearization measurements, a central wavelength of 550 nm was set in the power meter settings. Additionally, a spot luminance meter (LS-110, Konica Minolta, Inc., Tokyo, Japan) with a close-up lens was used to ensure the measured range and to link the measured light power values with actual luminance values.^21^

#### Software

The indoor-outdoor transition simulation was set up as a Unity3D scene using separated c#-scripts to control the different features of the testbed (Figure 5). To ensure that the pixel values of the walls can be set directly and are not influenced by light effects, the shader of the wall’s material was set to “unlit” from the universal render pipeline, rendering the objects, or here the walls, without any lighting effects. This way the overall VR room lighting could be controlled by accessing and modifying the wall’s material color specifications. The electrochromic lenses were simulated with a UI canvas attached to the main camera at a 0.1 m virtual distance. The specified transmission was achieved by accessing and modifying the canvas’ alpha specifications. For the conversion between transmission and alpha values the project’s color space was set to “gamma”.^21^ Additionally to the implications mentioned by Murray et al., clipping artifacts were observed when set to “linear” color space, where dark grey values were rendered as black thereby ruling out a realistic transmission simulation.

**Figure 5:**
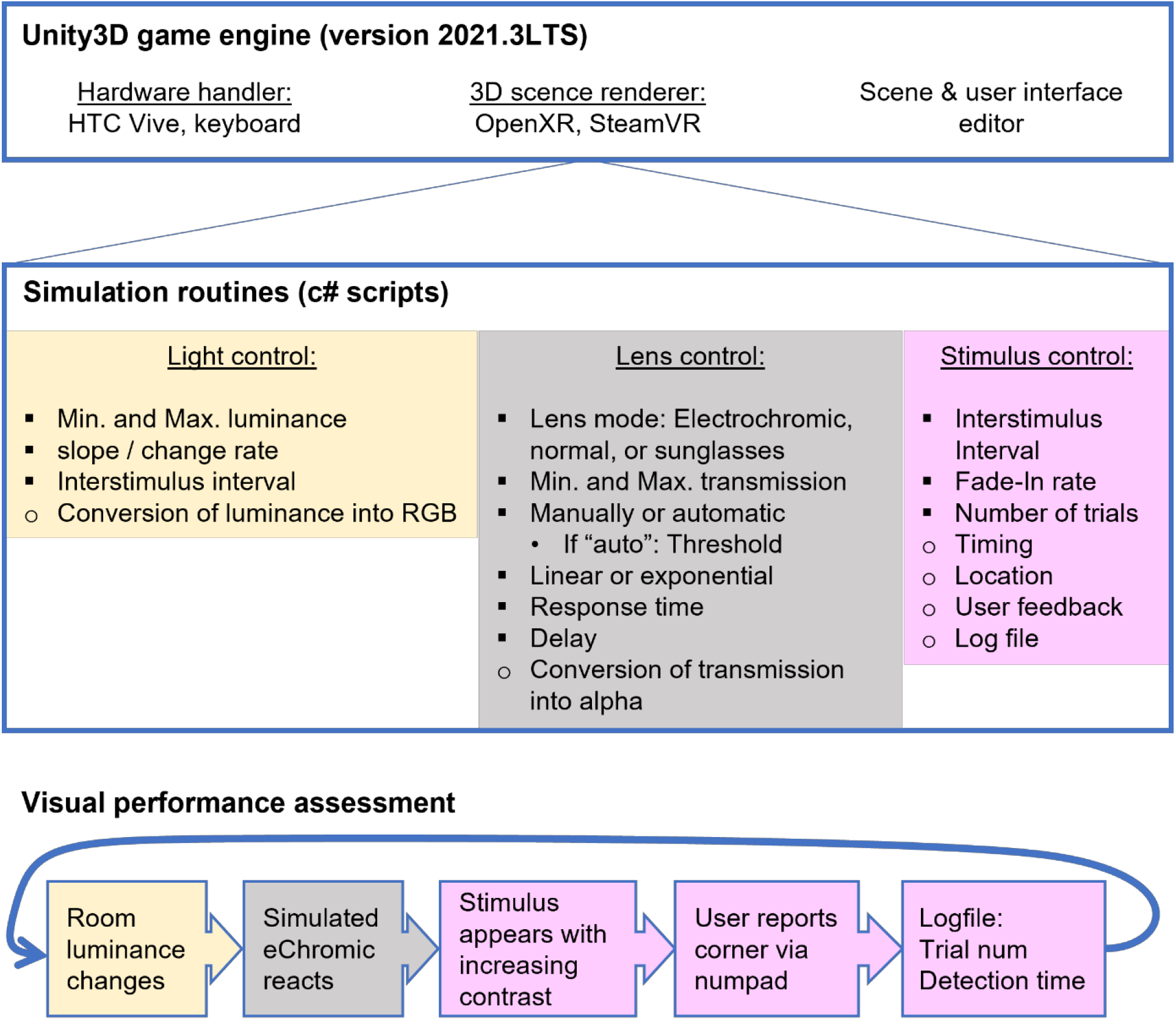
Basic software architecture and psychophysical assessment logic of the proposed testbed. Public variables that can be modified through the unity interface by an operator are marked with filled squares. Open circles denote routines embedded into the c#-scripts that cannot be modified externally.

The VR presentation of the unity scene on the HMD was enabled through the OpenXR packages in combination with the SteamVR (Valve Cooperation, Bellevue, WA, USA) plug-in.

### Psychophysical procedure

The general task for the participants was to detect a small achromatic low-contrast blob stimulus, appearing in one of four corners (top left, top right, bottom left, bottom right) at a 12 meter virtual distance, and report the correct corner via pressing the corresponding key on a numpad (Figure 2). The four possible stimulus locations were displaced from a central fixation target by about 13 degrees. The FWHM of the stimulus was about 0.6 degree. The stimulus material used the “unlit” shader of the universal render pipeline, and the stimulus contrast was controlled by adjusting the material’s color specifications.

To assess the impact of electrochromic lenses on visual performance following rapid luminance changes, participants had to detect the stimuli in eight different scenarios within 20 trials for room lights switching off and 20 trials for switching on, resulting in 40 trials in total per condition (see Table 2). The room light and the linked luminance were changed to simulate indoor-outdoor transitions. For the electrochromic lens, the response time (reaching 90 % of the target transmission value) was 3 or 1 seconds. Additionally, normal glasses with static 95 % transmission and sunglasses with static 20 % transmission were tested.

Participants underwent two training runs prior to the actual testing runs. The different conditions were tested in randomized order to minimize any influence from further training of the task during the ongoing experiment.

### Quantification and statistical analysis

The statistical analysis of the recorded detection times was done in MATLAB (The MathWorks, Inc., Natick, USA). First, outliers were identified as being more than 1.5 times the interquartile range higher than the 3rd quartile or lower than the 1st quartile and were removed. On average approximately 0.7 outliers per participant per condition occurred (minimum 0 – maximum 5 times), with a total of 115 outliers out of all 3200 trials recorded (3.6 %). Second, the participant’s median detection times were compared using the Wilcoxon signed rank test (Matlab: “signrank”), as data sets were not normally distributed (Kolmogorov-Smirnov test; Matlab: “kstest”).

